## Supplementary figure 1 and 2 for "Contrasting roles for IKK regulated inflammatory signalling pathways for development and maintenance of type 1 and adaptive γδ T cells"

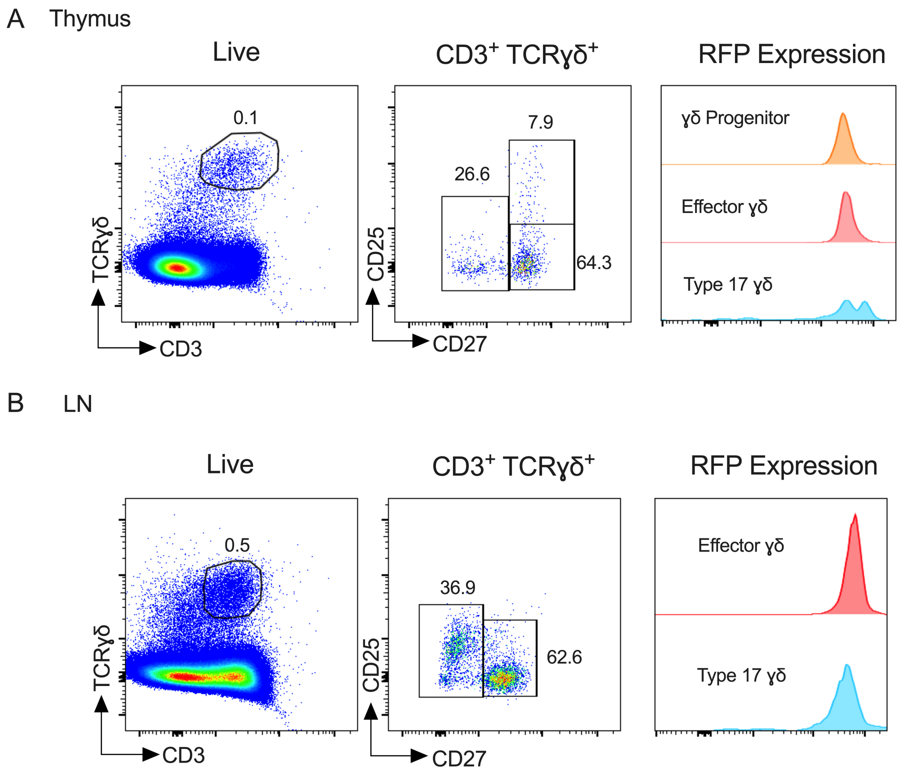


**Figure S1 - huCD2iCre targets Cre activity to γδ T cells in the thymus**

Thymi and lymph nodes (LNs) from Cre –ve mouse was analysed by flow. The density plot number represents the percentage of cells in each gate. (A) Representative flow plots are of thymocytes profiles: Histograms display RFP expression by γδ progenitor, effector γδ and type 17 γδ T cells. (B) Representative flow plots display LNs profiles: Histograms display RFP expression by effector γδ and type 17 γδ T cells.

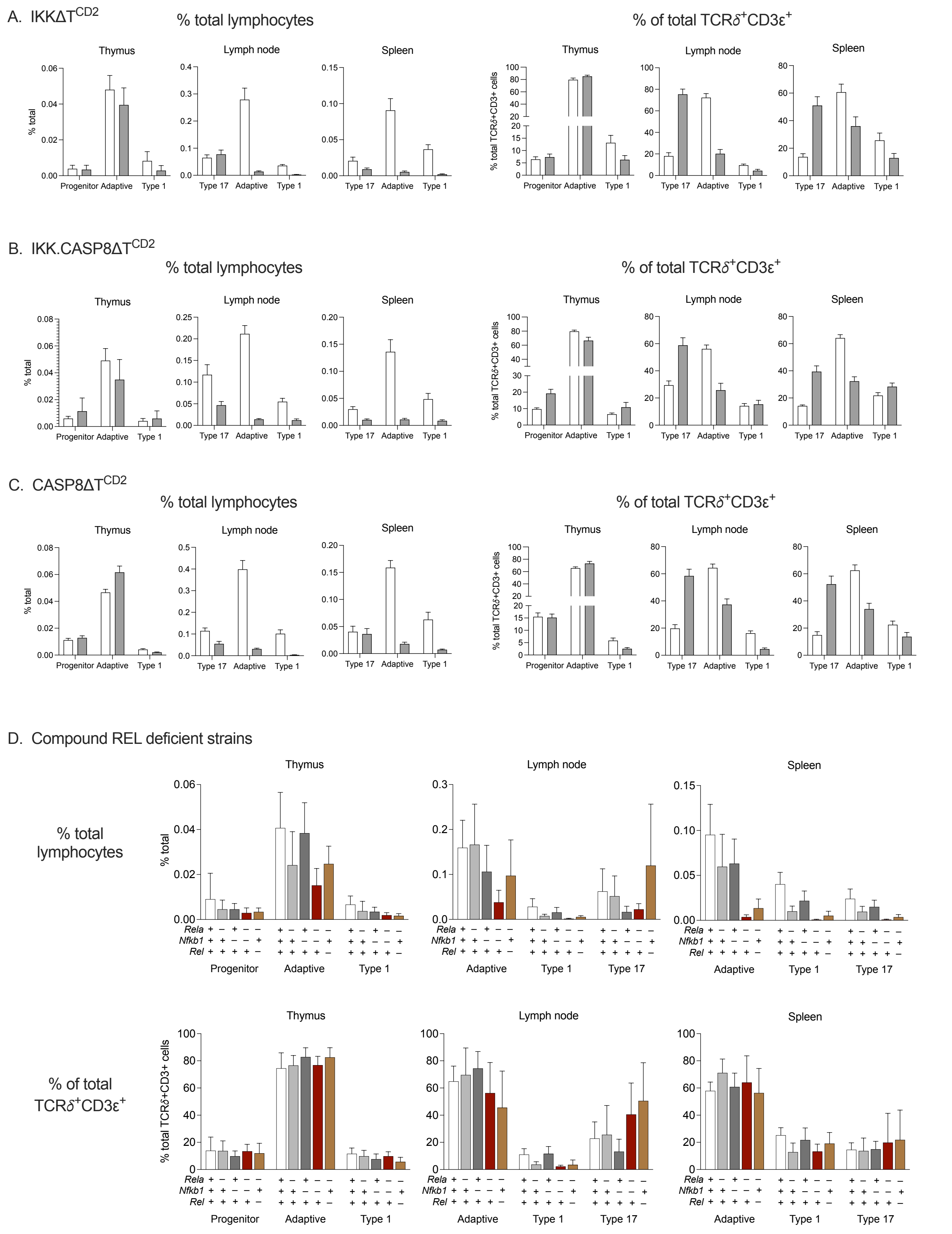


**Figure S2 - Representation of different ɣδ T cell sub-populations in different mouse strains.** Bar charts show the representation of progenitor, adaptive and type 1 ɣδ T cell subsets in the thymus, and adaptive, type 1 and type 17 subsets in lymph nodes and spleen. Charts are either of percentage representation amongst total lymphocytes of the specified organ (% total lymphocytes), or the relative representation of the indicated subset amongst total gd T cells in the given organ (% of total TCR$\delta^{+}$ CD3$\varepsilon^{+}$). Data are shown for IKKΔT$^{CD2}$ mice described in figure 1, Casp8.IKKΔT$^{CD2}$ mice described in figure 2, Casp8ΔT$^{CD2}$ mice described in figure 4 and RelaΔT$^{CD2}$, *Nfkb1*$^{-/-}$, RelaΔT$^{CD2}$ *Nfkb1*$^{-/-}$ mice, Rela.RelΔT$^{CD2}$ mice described in figure 7.
